# Martini 3 Coarse-Grained Model of DNA for Heterogeneous Molecular Systems

**DOI:** 10.64898/2026.09.14.751576

**Authors:** Rokas Dargis, Gaurav Arya

**Affiliations:** Department of Mechanical Engineering and Materials Science, Duke University, Durham

**Keywords:** Martini 3 coarse-grained model, molecular dynamics simulations, DNA, Bayesian optimization, Wasserstein distance, DNA nanotechnology

## Abstract

DNA often functions in heterogeneous molecular systems containing proteins, lipids, polymers, and other materials. All-atom molecular dynamics simulations can be used to study DNA in these multicomponent systems, but computational cost limits the accessible system sizes and time scales. Coarse-grained models extend these scales, but existing DNA models are generally not designed for interactions with a broad range of other molecular species. To fill this gap, we develop a coarse-grained model of DNA designed for use with the Martini 3 force field. The model was parameterized through an iterative Bayesian optimization workflow, which used a scaled Wasserstein metric to compare distributions of local geometrical features and global structure from coarse-grained simulations against all-atom reference simulations. The optimized model captures key structural and mechanical properties of single- and double-stranded DNA across varying strand lengths and ionic conditions, while retaining compatibility with the broader Martini 3 ecosystem. This compatibility enables DNA to be integrated with a broad range of molecular systems, as we illustrate through simulations of double-stranded DNA bound to a transcription factor, cholesterol-tagged DNA duplex interacting with a lipid bilayer, a crossover-containing DNA nanostructure, and single-stranded DNA adsorbing onto graphene. Together, these results establish a transferable coarse-grained model of DNA for simulations of heterogeneous biomolecular and engineered systems.

## Introduction

DNA participates in chemically diverse systems involving a broad range of molecular interactions, enabled by its structural and electrostatic properties [1, 2]. These interactions underlie processes such as protein recognition, genome organization, and DNA packaging [3–5], and are increasingly exploited in engineered systems including DNA origami, nucleic-acid delivery platforms, and membrane-associated DNA assemblies [6–8]. Simulating such systems therefore requires models that can capture DNA and its interactions with other molecular species in chemically heterogeneous environments.

All-atom (AA) molecular dynamics (MD) simulations can describe the structure and interactions of DNA in molecular detail [1], but their computational cost limits the length and time scales accessible for large DNA-containing assemblies [9]. Coarse-grained (CG) DNA models address this limitation and have enabled simulations of DNA mechanics, hybridization, and DNA nanotechnology [9]. For example, oxDNA has been highly successful for modeling sequence-dependent DNA behavior and programmable DNA assemblies [10, 11], but it is primarily designed to model nucleic-acids rather than for integration within a broader biomolecular and soft-matter force-field ecosystem.

The Martini force field provides a general CG framework for modeling biomolecular and materials systems using a limited set of interaction sites that represent groups of atoms and are parameterized to reproduce structural and thermodynamic properties [12]. This common representation allows chemically distinct molecular species to be described within the same interaction framework. The original Martini force field was developed primarily for lipid systems [13] and was subsequently expanded in Martini 2 to encompass a much broader range of biomolecules [12]. A CG DNA model was also developed within Martini 2, enabling simulations of DNA together with other components represented by the force field [14]. Martini 3 further expands the chemical scope and transferability of the framework, with established models for proteins, lipids, carbohydrates, and RNA [15–17], as well as synthetic material systems such as polymers and carbon nano-materials [18, 19]. Extending DNA to Martini 3 would therefore enable its integration with a broader range of molecular species and facilitate simulations of chemically diverse biomolecular and engineered systems.

Here, we developed such a coarse-grained model for DNA using existing Martini 3 bead types and the standard non-bonded interaction framework, reducing the need to parameterize new pairwise interactions for each mixed system [15]. Backbone bonded parameters were optimized against all-atom reference simulations using an iterative Bayesian optimization (BO) workflow, with a scaled Wasserstein metric comparing local backbone and global duplex structural distributions between CG and AA trajectories 1. The resulting model was validated and then evaluated across multiple DNA-containing systems designed to test key applications enabled by Martini 3 compatibility: double-stranded DNA (dsDNA) bound to the *λ* repressor protein, cholesterol-tagged dsDNA associating with a lipid bilayer, a crossover-containing DNA nanostructure, and single-stranded DNA (ssDNA) adsorption onto graphene. Across all systems, the model maintained structurally stable DNA conformations, supporting its use for Martini 3 simulations of DNA in heterogeneous biomolecular and soft-matter environments.

## Methods

### Coarse-Grained Representation of DNA

The Martini 3 DNA model was constructed from the existing Martini 3 RNA representation [17] through two main developments: redefinition of the coarse-grained mapping to represent DNA chemistry and reparameterization of the backbone bonded interactions for dsDNA. The RNA mapping was retained where possible, with two modifications to represent the chemical differences between RNA and DNA: the uracil mapping was modified to represent thymine, and the BB3 backbone bead was redefined to exclude the 2^′^-hydroxyl oxygen present in ribose but absent in deoxyribose [1, 20]. To account for differences in the phosphate backbone of RNA and DNA, the backbone bonded parameters were selected for optimization against an all-atom dsDNA reference to reproduce DNA-specific backbone geometry. These comprised the equilibrium value and force constants of the three bond lengths, three angles, and three proper dihedrals between backbone beads, as depicted in Figure 2A. This resulted in a total of 18 parameters to be optimized. Other bonded interactions were inherited from the Martini 3 RNA model and retained without modification, including interactions within the side chains and between side-chain and backbone beads. The fixed elastic network used in Martini 2 DNA to maintain the duplex structure was also retained[14].

### Bayesian Optimization of Backbone Parameters

The 18 backbone bonded parameters described above were optimized using an iterative BO pipeline (Fig. 1) implemented in Python with the *bayes_opt* package [21]. The optimizer proposed backbone parameter sets within predefined bounds (Table S1), which were used to carry out a new CG MD simulation. The simulated CG ensemble was then evaluated against a AA reference. Based on the evaluation score, new parameter sets were proposed using a Gaussian-process surrogate model [22] with a stochastic mean–uncertainty acquisition function. For each suggestion, 10,000 candidate parameter sets were sampled uniformly, and the Gaussian-process posterior mean, *µ*(*x*), and standard deviation, *σ*(*x*), were evaluated for each candidate. A scalar *β* was independently sampled from a standard normal distribution, and candidates were ranked according to:

**Figure 1:**
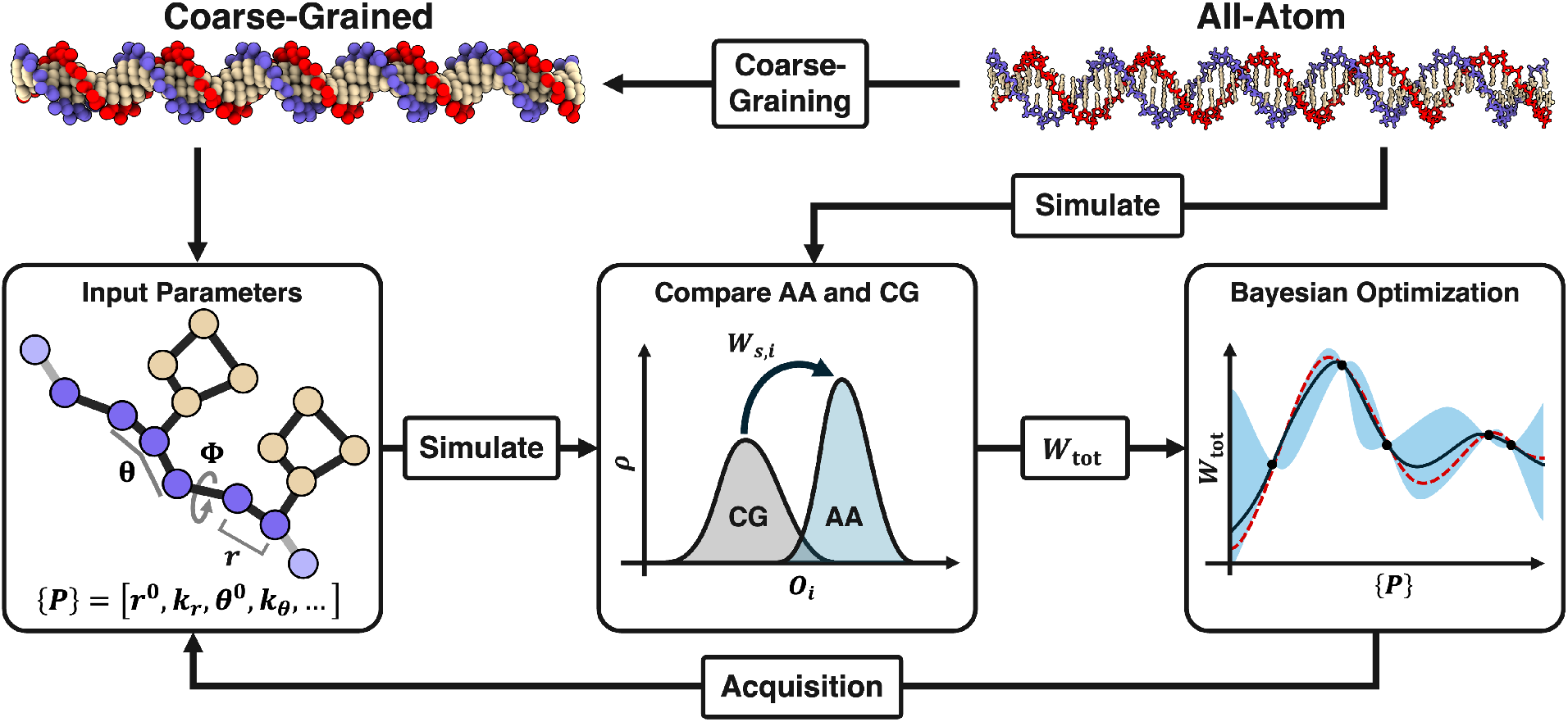
Iterative Bayesian optimization pipeline for parameterizing the coarse-grained dsDNA model. The workflow begins by coarse-graining an AA dsDNA structure to generate the initial CG representation. The AA structure is simulated using a fixed AA parameter set, and the resulting trajectory serves as the target for optimization. In each BO iteration, the CG model is simulated using a proposed backbone parameter set {*P*} containing the equilibrium values and force constants for backbone bonds (*r*^0^, *k*_*r*_), angles (*θ*^0^, *k*_*θ*_), and dihedrals (*ϕ*^0^, *k*_*ϕ*_). Structural observables *O*_*i*_ are extracted from both AA and CG trajectories, and their distributions are compared using the scaled Wasserstein metric, *W*_s,*i*_. The individual *W*_s,*i*_ values are summed into *W*_tot_, which is returned to the optimizer to update the Gaussian-process surrogate model and guide the selection of the next parameter sets.

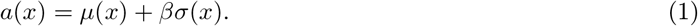

The candidate with the highest acquisition value was selected for simulation. A new value of *β* was drawn for each suggestion, allowing the relative contribution of the posterior mean and uncertainty to vary between suggestions. Parameter sets were generated in batches of 20 and simulated in parallel, with the resulting scores registered with the optimizer after completion of each batch. A total of 20 batches were evaluated. The first 10 batches used the Gaussian-process acquisition procedure exclusively. Mutation batches were then introduced at batches 11, 15, and 19, while Gaussian-process acquisition was for the intervening batches and the final batch.

For each mutation batch, mutation began from the parameter set with the best overall evaluation score observed at that stage of optimization. Each of the 18 parameters had a 50% probability of being retained from the current best parameter set and a 50% probability of being replaced. Replacement values were randomly drawn from a candidate pool containing the corresponding parameter values from the next four best-performing parameter sets together with the parameter value that produced the best individual score for the corresponding local observable. Mutated parameter sets were then simulated and evaluated using the same procedure as acquisition-generated parameter sets.

### Structural Observables and Model Scoring

The parameter sets generated during each optimization batch were used to carry out 100 ns long CG simulations of a 50 bp B-DNA duplex of the following randomly generated sequence: 5^′^-AAGTTTAACAA ATTATCTCGATCGGTTGAACAGGGACACAGGTAATGAGG-3^′^. Each parameter set was evaluated by comparing the structural distributions of its CG trajectory to an 100 ns long AA reference simulation of the same sequence. To enable direct comparison, the AA trajectory was mapped onto the same coarse-grained representation used for the CG model before calculating the structural distributions.

We evaluated each CG parameter set by comparing distributions of local and global structural observables from the CG trajectory with those from the AA trajectory. Local observables characterized the optimized backbone geometry by measuring the bond lengths, angles, and dihedrals between backbone beads: *O*_L_ = {*r*_1_, *r*_2_, *r*_3_, *θ*_1_, *θ*_2_, *θ*_3_, *ϕ*_1_, *ϕ*_2_, *ϕ*_3_} (Fig. 2A). Global observables characterized the duplex structure through the terminal distance, minor and major groove widths, and backbone separation between complementary nucleotides: *O*_G_ ={*d*_term_, *d*_minor_, *d*_major_, *d*_bb_} (Fig. 2B).

**Figure 2:**
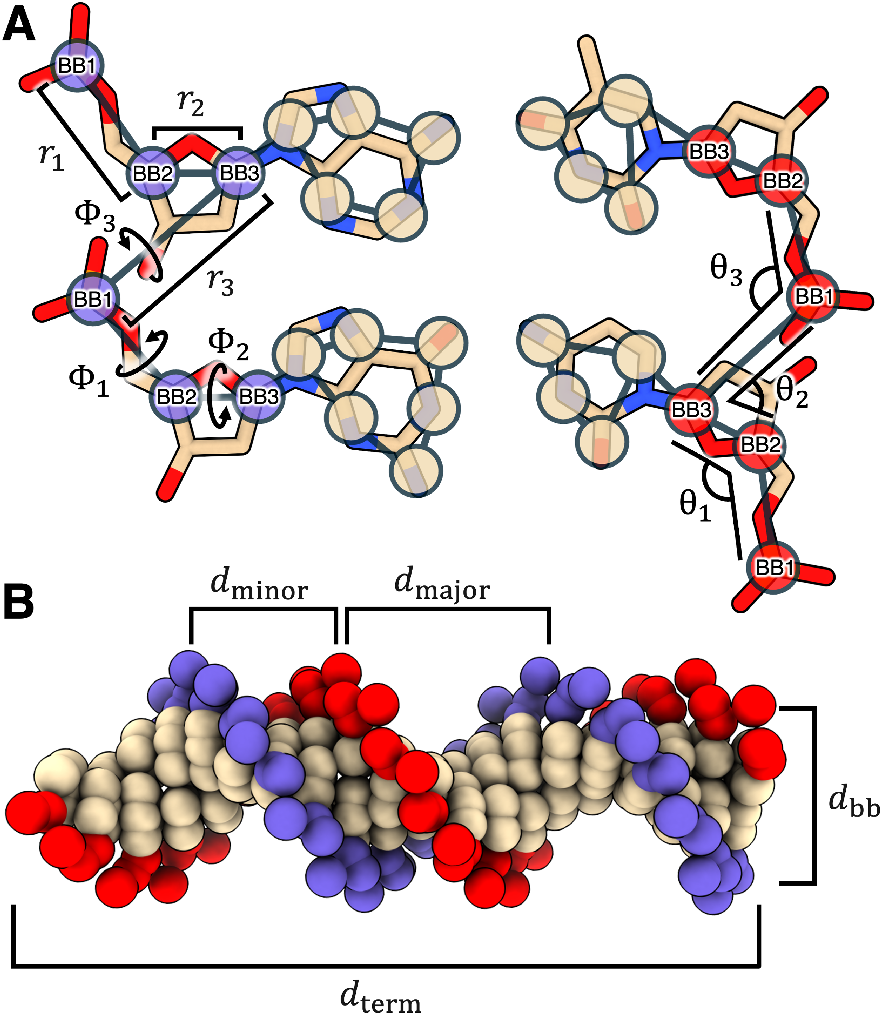
Backbone bonded terms and structural observables used for Martini 3 DNA parameterization. (A) Three backbone bonds (*r*_1_, *r*_2_, *r*_3_), three angles (*θ*_1_, *θ*_2_, *θ*_3_), and three dihedrals (*ϕ*_1_, *ϕ*_2_, *ϕ*_3_) define the bonded interactions whose equilibrium values and force constants were optimized during parameterization. These same bond lengths, angles, and dihedrals were used as the local structural observables for evaluating each parameter set. (B) Global observables describe the structure of dsDNA and include the terminal distance (*d*_term_), minor and major groove widths (*d*_minor_, *d*_major_), and backbone separation between paired nucleotides (*d*_bb_).

To quantify how closely the CG distributions matched the AA reference distributions, we used a scaled form of the Wasserstein metric. The standard one-dimensional Wasserstein metric *W* computes the optimal transportation cost required to transform one distribution into another by moving probability mass along the observable coordinate [23, 24]. For each observable *i*, the metric *W*_*i*_ was computed between the AA and CG cumulative distribution functions:

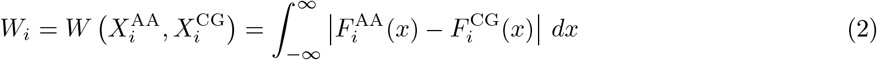

where 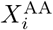 and 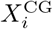 are the sampled AA and CG distributions for observable *i*, 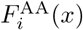 and 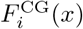 are their corresponding cumulative distribution functions, and *x* is the value of observable *i* over which the distributions are integrated. Because *W*_*i*_ retains the units and scale of the underlying observable, it cannot be directly compared across observables with different units and magnitudes. Each *W*_*i*_ was therefore normalized by an observable-specific scale factor derived from the AA reference distribution:

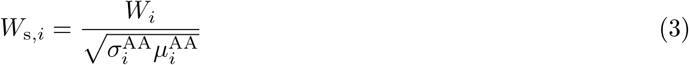

where 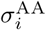 and 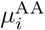 are the standard deviation and mean of the AA reference distribution for observable *i*, respectively. For each parameter set, the scaled Wasserstein values were combined into a single loss *W*_tot_ according to:

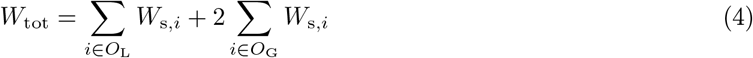

The global contribution was weighted by a factor of two to reduce the imbalance between the four global and nine local observables and prevent the local observable set from dominating the optimization objective. Lower *W*_tot_ values indicate closer agreement between the CG and AA structural distributions. Because the BO implementation maximizes its objective function, −*W*_tot_ was registered with the optimizer, such that maximizing the BO objective was equivalent to minimizing the AA–CG structural discrepancy.

### Simulation Methods

All CG simulations were performed using GROMACS [25]. The Martini 3 force field [15] was used throughout, with the exception of the Martini 2 comparison simulations [12]. Unless otherwise specified, systems were neutralized with Na^+^ counterions and supplemented with NaCl to a concentration of approximately 150 mM. Systems were energy minimized by steepest descent, followed by 0.5 ns of NVT equilibration and three NPT equilibration stages of 2, 10, and 10 ns. Position restraints were applied during equilibration and removed for production unless otherwise specified. Temperature and pressure were maintained at 310 K and 1 bar using the velocity-rescale thermostat and C-rescale barostat, respectively. Reaction-field electrostatics were used with a relative dielectric constant of 15 and a cutoff of 1.1 nm, and van der Waals interactions used a 1.1 nm cutoff with the potential-shift-Verlet modifier [15, 26]. Production simulations used a 5 fs timestep unless otherwise specified. The initial portion of each production trajectory was excluded from analysis to allow relaxation from the starting configuration.

The AA reference simulation was performed using AMBER22 [27] with the parmbsc1 DNA force field [28] and OPC water model [29]. The system was solvated in a truncated-octahedral water box, neutralized with Na^+^ counterions, and supplemented with NaCl to approximately 150 mM. Following energy minimization and heating to 310 K, production was performed at 310 K and 1 bar using a 2 fs timestep. Bonds involving hydrogen were constrained using SHAKE [30], long-range electrostatics were treated using particle-mesh Ewald [31], and a 0.9 nm cutoff was used for short-range non-bonded interactions.

System-specific details, including DNA sequence and length, ion concentrations, production lengths, and deviations from these protocols, are summarized in the Supporting Information (Tables S2 and S3).

## Results

### Parameter Optimization

The BO workflow progressively identified parameter sets with improved AA–CG agreement (Fig. 3A). While the running batch average decreased gradually over the course of optimization, later batches continued to identify parameter sets with lower *W*_tot_. In particular, the first mutation batch at batch 11 produced a marked decrease in the minimum *W*_tot_. Additional acquisition and mutation batches were subsequently evaluated, but no lower *W*_tot_ was identified after batch 11. The lowest-scoring parameter set observed across the optimization was therefore selected as the final model. The resulting optimized parameters are reported in Table 1, and distributions of the sampled structural observables for the final model are shown in Fig. S1. As *W*_tot_ improved, most individual parameters shifted toward regions associated with lower local *W*_s_ values (Fig. 3B). However, the final parameter set did not coincide with the lowest local *W*_s_ regions for every parameter, indicating that minimizing the combined objective required balancing agreement across multiple local and global observables.

**Table 1:** Optimized Martini 3 DNA backbone bonded parameters.

| Equilibrium value |  | Force constant |  |
| --- | --- | --- | --- |
| $r_1^0$ | 0.351 nm | $k_{r_1}$ | 5630 kJ/(mol nm<sup>2</sup>) |
| $r_2^0$ | 0.176 nm | $k_{r_2}$ | 21900 kJ/(mol nm<sup>2</sup>) |
| $r_3^0$ | 0.390 nm | $k_{r_3}$ | 15000 kJ/(mol nm<sup>2</sup>) |
| $\theta_1^0$ | 112° | $k_{\theta_1}$ | 799 kJ/mol |
| $\theta_2^0$ | 80.1° | $k_{\theta_2}$ | 529 kJ/mol |
| $\theta_3^0$ | 75.8° | $k_{\theta_3}$ | 979 kJ/mol |
| $\phi_1^0$ | 140° | $k_{\phi_1}$ | 4.97 kJ/mol |
| $\phi_2^0$ | 163° | $k_{\phi_2}$ | 2.01 kJ/mol |
| $\phi_3^0$ | 15.4° | $k_{\phi_3}$ | 21.3 kJ/mol |

**Figure 3:**
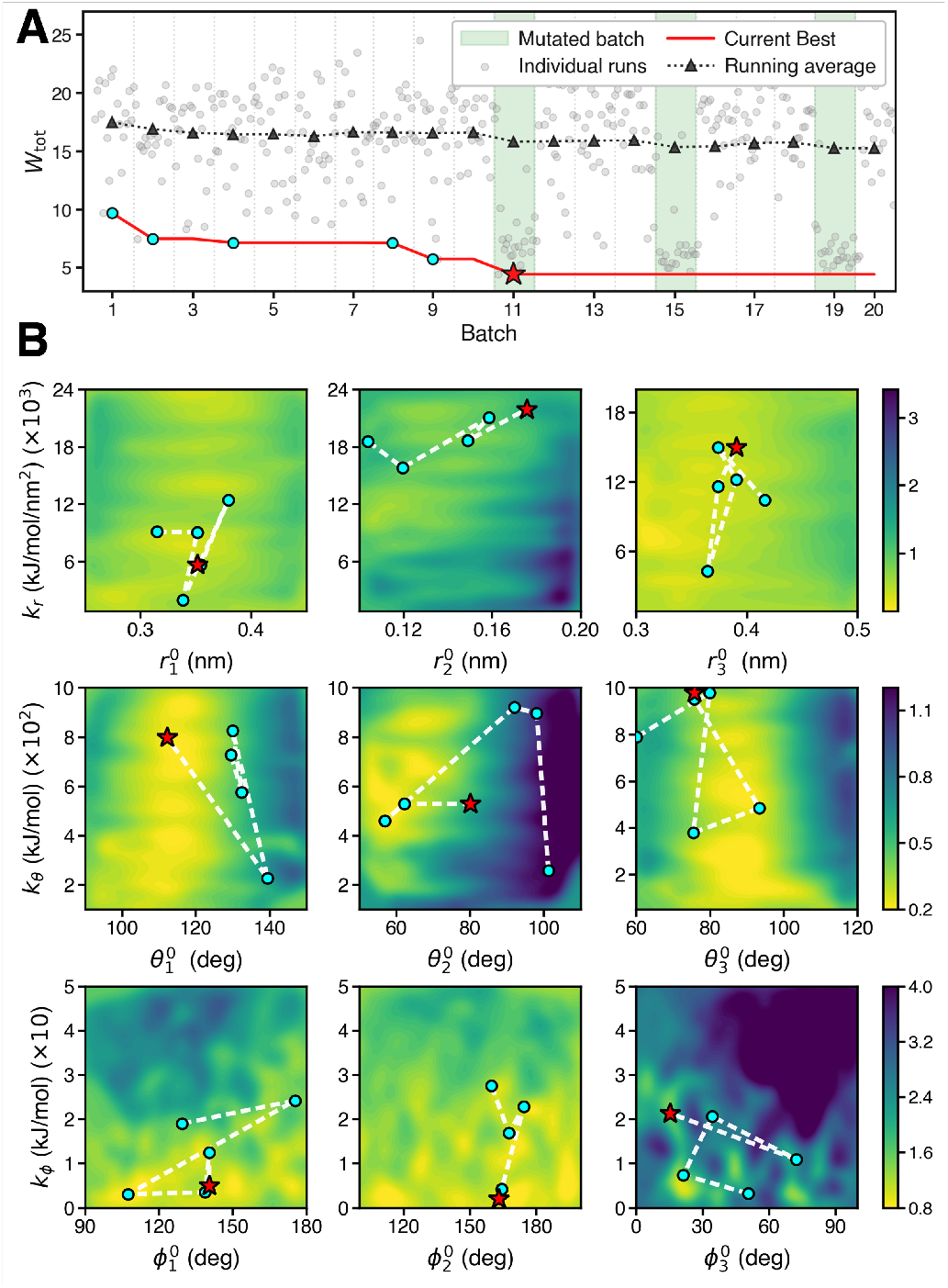
Optimization trajectory of the CG dsDNA backbone parameters. (A) Bayesian optimization across 20 batches using Gaussian-process acquisition (white background) and parameter mutation (green background). Gray points show *W*_tot_ for each parameter set, black triangles show the batch average, and the red line indicates the lowest *W*_tot_ observed up to each batch. Cyan circles mark successive improvements in the lowest *W*_tot_, and the red star indicates the final selected parameter set. (B) *W*_s_ landscape across equilibrium value-force constant pairs, generated by interpolating and smoothing the sampled scores. Color bars indicate the corresponding *W*_s_ values. Cyan points trace parameter values corresponding to successive improvements in the best *W*_tot_, and red stars mark the final selected parameter set, corresponding to panel A.

As an additional assessment of the scoring metric, we compared the scaled Wasserstein score with Kullback-Leibler divergence (*D*_KL_), which is commonly used for comparing distributions [32]. Under limitedoverlap conditions, *D*_KL_ can become very large or formally divergent when one distribution assigns little or no probability to regions sampled by the other, whereas the Wasserstein distance remains well defined for distributions with limited-overlap [33]. This limitation is particularly relevant for BO, where exploratory parameter sets often produced CG distributions that were substantially shifted from the AA reference. The Wasserstein metric, in contrast, remained finite and continued to quantify the separation between distributions even when their overlap was small. Visual comparison of representative AA and CG distributions showed that the scaled Wasserstein metric more consistently tracked the apparent degree of distributional agreement than Kullback-Leibler divergence (Fig. S2).

### Validation of DNA Model

To evaluate the accuracy and transferability of the optimized model, we performed additional DNA simulations spanning different lengths, sequences, and ionic conditions. These systems were compared against Martini 2, all-atom force fields, and available experimental measurements.

We first benchmarked the Martini 3 DNA model against the Martini 2 DNA model using both soft and stiff elastic networks [14]. All three models were evaluated using the same local and global observables (Fig. 2) against the same AA reference distributions, allowing direct comparison of their overall agreement with the atomistic reference. Overall, the Martini 3 model achieved the lowest *W*_tot_, with a value of 4.41 compared with 12.46 and 8.54 for the soft and stiff Martini 2 models, respectively (Fig. 4A). For 9 of the 13 individual observables, Martini 3 produced the lowest *W*_s_. These improvements spanned both local geometrical features and global duplex structure, indicating that the Martini 3 model showed closer overall agreement with the AA reference than either Martini 2 representation across the structural metrics evaluated here.

**Figure 4:**
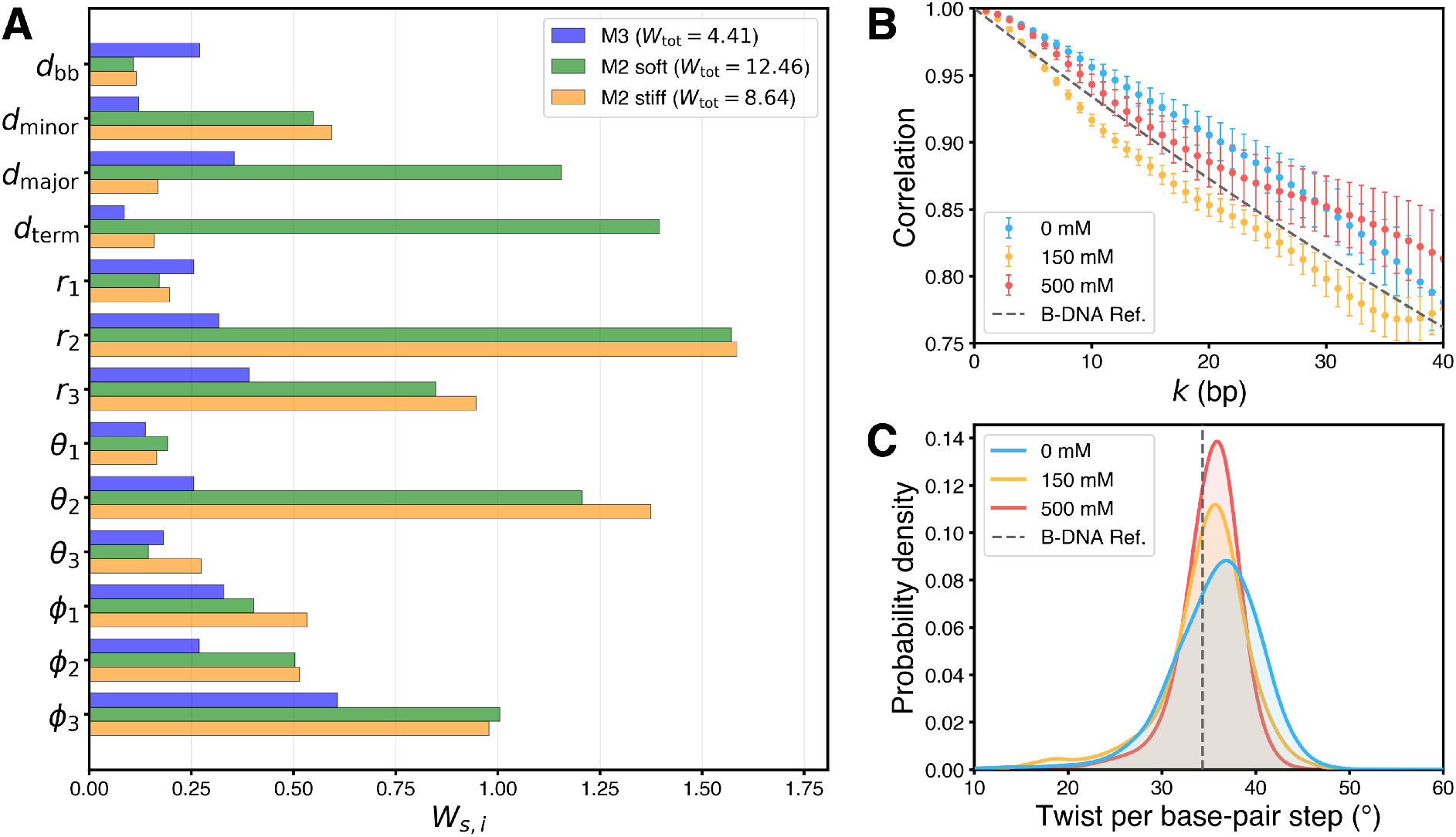
Structural evaluation of the optimized Martini 3 dsDNA model. (A) Comparison of scaled Wasserstein scores *W*_s_ for each observable between the optimized Martini 3 model (blue), and the Martini 2 model using soft (green) and stiff (yellow) elastic networks. (B) Tangent correlation as a function of base-pair separation, *k*, for 100 bp dsDNA under 0 mM (green), 150 mM (purple), and 500 mM (red) ionic conditions. Error bars represent the error of the mean of the correlation across three time blocks of the trajectory. Gray dashed line indicates the expected correlation of dsDNA with 50 nm persistence length. (C) Twist per base-pair step twist distributions at the same three ionic conditions. Gray dashed line indicates the canonical 34.3° twist of B-DNA.

Second, we assessed the persistence length and base-pair step twist of 100 bp dsDNA at three added-salt concentrations: 0, 150, and 500 mM NaCl, with neutralizing counterions present in all systems. DNA flexibility was evaluated from the tangent correlation along the duplex axis:

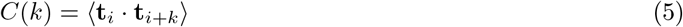

where **t**_*i*_ and **t**_*i*+*k*_ are local tangent vectors separated by *k* base pairs. Local tangent vectors were defined from the displacement between base-pair centers separated by 10 bp, approximately one helical turn, to suppress oscillations arising from individual base-pair centers not lying on the helical axis (Fig. 4B). The persistence length *L*_*p*_ was obtained by fitting the tangent correlation to the worm-like chain model [10]:

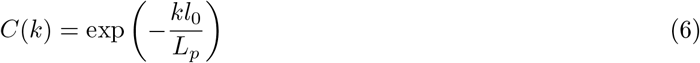

where *l*_0_ is the mean rise per base pair. The resulting persistence lengths were 69.1 ± 10.2 nm, 51.3 ± 3.8 nm, and 78.8 ± 16.8 nm for 0 mM, 150 mM, and 500 mM systems, respectively, where the uncertainties represent the standard error of mean across persistence lengths independently fitted to five blocks of the analyzed trajectory. Although no clear salt-dependent trend was resolved, consistent with the Martini 2 soft-network results, the persistence lengths remained around 50 nm, broadly matching both the Martini 2 soft-network values and the experimental persistence length of approximately 45–50 nm at moderate-to-high ionic strength [14, 34].

Helical structure was independently evaluated from the twist between consecutive base pairs. For each base pair, a local coordinate frame was constructed from the backbone and nucleobase beads, and the rotation between adjacent base-pair frames about the local helical axis was calculated to obtain the base-pair step twist. The corresponding helical repeats were 10.36 ± 0.05, 10.61 ± 0.08, and 10.52 ± 0.02 bp per turn for the 0 mM, 150 mM, and 500 mM systems, respectively (Fig. 4C), closely matching the canonical B-DNA value of approximately 10.5 bp per turn [20]. Together, these results show that the model maintains B-DNA-like helicity while capturing long-range duplex stiffness not explicitly included as targets during parameterization.

Lastly, to assess how well the optimized backbone parameters for dsDNA reproduce ssDNA structure in solution, we simulated ssDNA strands ranging from 10 to 50 nucleotides at 12.5 mM and 125 mM ionic strength and calculated their radius of gyration (*R*_*g*_):

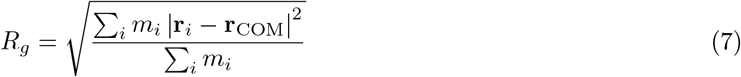

where *m*_*i*_ and **r**_*i*_ are the mass and position of bead *i*, respectively, and **r**_COM_ is the center of mass of the ssDNA strand. The Martini 3 results were compared with previously reported Martini 2, CHARMM, and AMBER simulations performed under counterion-only and 100 mM NaCl conditions, together with experimental measurements of poly-dA and poly-dT at 12.5 mM and 125 mM ionic strength [14, 35]. Across strand lengths, Martini 3 followed the trends of the previously reported computational models. Compared with the experimental measurements, Martini 3 produced more compact ssDNA conformations, consistent with the discrepancy between simulated and experimental *R*_*g*_ values previously observed for Martini 2 and the atomistic models (Fig. 5). An additional concentration of 1205 mM, substantially above physiological ionic strength, was tested to examine ssDNA behavior at very high salt concentration (Fig. S3).

**Figure 5:**
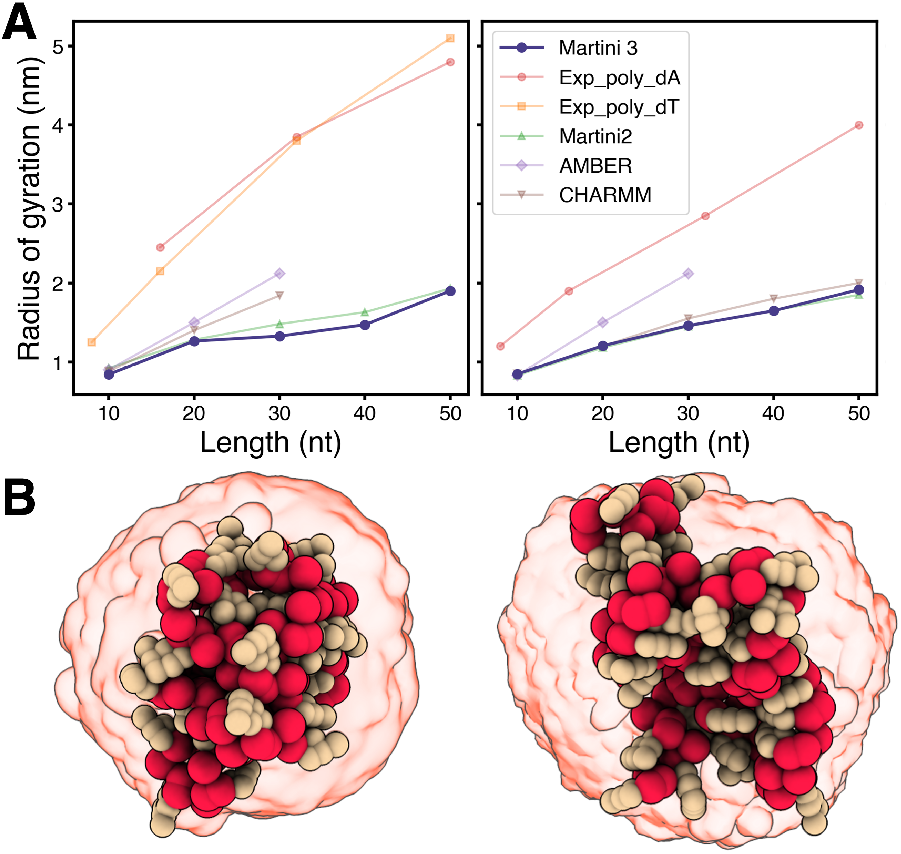
Validation of ssDNA radius of gyration. (A) Radius of gyration as a function of ssDNA strand length for the optimized Martini 3 model, Martini 2, AMBER, CHARMM, and experimental SAXS measurements for poly-dA and poly-dT [14, 35]. Martini 3 and the experimental measurements correspond to 12.5 mM (left panel) and 125 mM ionic strength (right), while the previously reported Martini 2, AMBER, and CHARMM simulations correspond to counterion-only and 100 mM NaCl conditions, respectively. (B) Representative snapshots of a 40 nt Martini 3 ssDNA strand at each concentration. Beaded conformations show a single configuration at 90 ns, and the red transparent surface represents the conformational volume sampled across the trajectory.

### Applications in Multicomponent Systems

To evaluate the model beyond isolated DNA systems, we tested it across four distinct application environments spanning protein binding, membrane association, adsorption to a non-biological material, and crossover-containing DNA nanostructures. These systems were selected to assess whether the validated Martini 3 DNA model could maintain stable DNA conformation while interacting with independently parameterized Martini 3 biomolecular and material components.

To begin, we simulated a *λ* repressor protein bound to its dsDNA operator (PDB ID: 1LMB) [36] for 500 ns as a protein-DNA interaction application. Visual inspection of the trajectory showed that the repressor remained bound to DNA throughout the simulation, with no dissociation observed (Fig. 6A). The root-mean-square deviation (RMSD) of the dsDNA, calculated relative to the average structure over the trajectory, started at approximately 3.0 Å and stabilized at about 2.0 Å over the course of the simulation (Fig. 6B), which are comparable in magnitude to those of the same complex computed using the Martini 2 model [14].

**Figure 6:**
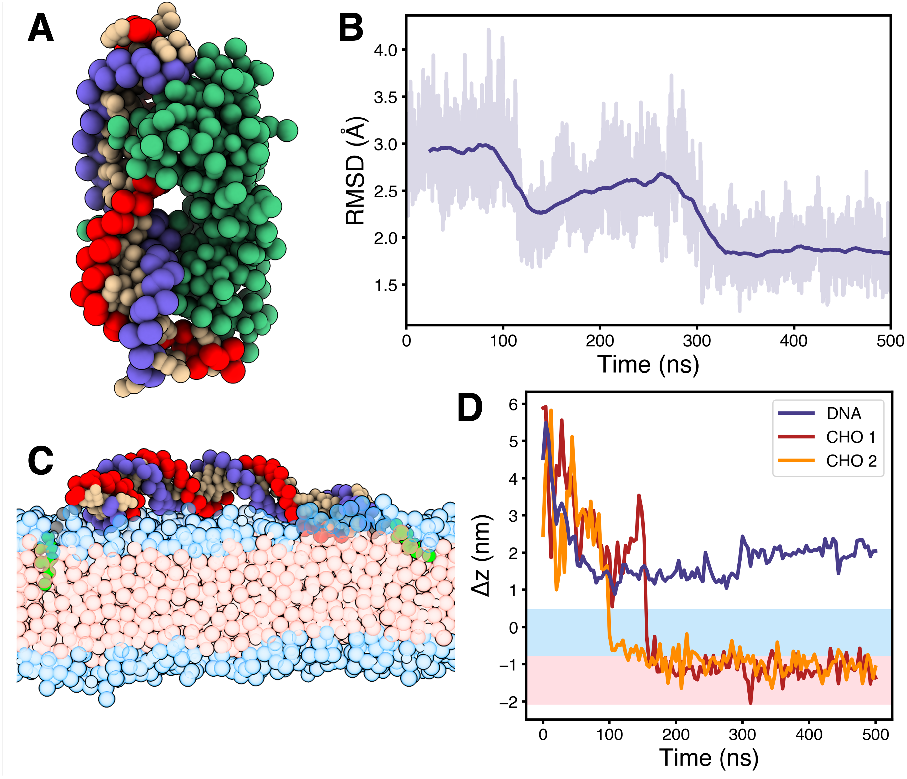
Martini 3 DNA model in protein- and lipid-associated systems. (A) Snapshot of coarse-grained dsDNA (blue and red backbones with tan nucleobases) bound to *λ* repressor (green) after 500 ns of simulation. (B) RMSD of the dsDNA in the protein complex relative to the average structure. The light trace shows the instantaneous RMSD and the dark trace a 20 ns rolling average. (C) Snapshot of cholesterol-tagged dsDNA associated with POPC bilayer after 200 ns of simulation. The cholesterol anchors (green) extend past the headgroup region (blue) and into the hydrophobic tail region (pink). (D) Center-of-mass positions of dsDNA (purple line) and cholesterol (orange and red lines) relative to the upper POPC leaflet along the membrane-normal. Shaded regions indicate the headgroup (blue) and hydrophobic tail (pink) regions.

To evaluate the model in a lipid bilayer-associated DNA system, we simulated cholesterol-tagged dsDNA interacting with a POPC bilayer [8, 37]. Cholesterol was attached to each 5^′^ end of a 30 bp DNA duplex through a six-carbon linker (Fig. S4) [38], and the system was simulated for 500 ns using the Martini 3 lipid model [39]. At approximately 100 ns, one cholesterol anchor inserted past the hydrophilic lipid headgroups into the hydrophobic bilayer core, followed shortly by the second anchor (Fig. 6C,D). Following insertion, the cholesterol anchors diffused laterally within the bilayer while the tethered dsDNA remained mobile, bending and occasionally separating from the surface without loss of duplex integrity.

As a non-biological material application, we simulated ssDNA starting above a graphene surface parameterized using the Martini 3 graphene model developed by Shrestha *et al*. [19]. Initially, the ssDNA remained mobile above the graphene surface, with the vertical center-of-mass separation between the ssDNA and graphene fluctuating by several nanometers (Fig. 7A,B). After approximately 50 ns, the strand rapidly approached and adsorbed onto the graphene, decreasing its vertical separation to approximately 1.3 nm. The ssDNA remained associated with the surface for the remainder of the 500 ns simulation and underwent further rearrangement on the surface, reaching a separation of approximately 1.0 nm after ∼180 ns. This spontaneous and persistent adsorption is consistent with previously reported ssDNA-graphene behavior [40]. Upon adsorption, some nucleobases adopted T-shaped *π*-*π* stacking orientations relative to the graphene surface (Fig. S5), consistent with previously reported nucleobase–graphene interactions [41].

**Figure 7:**
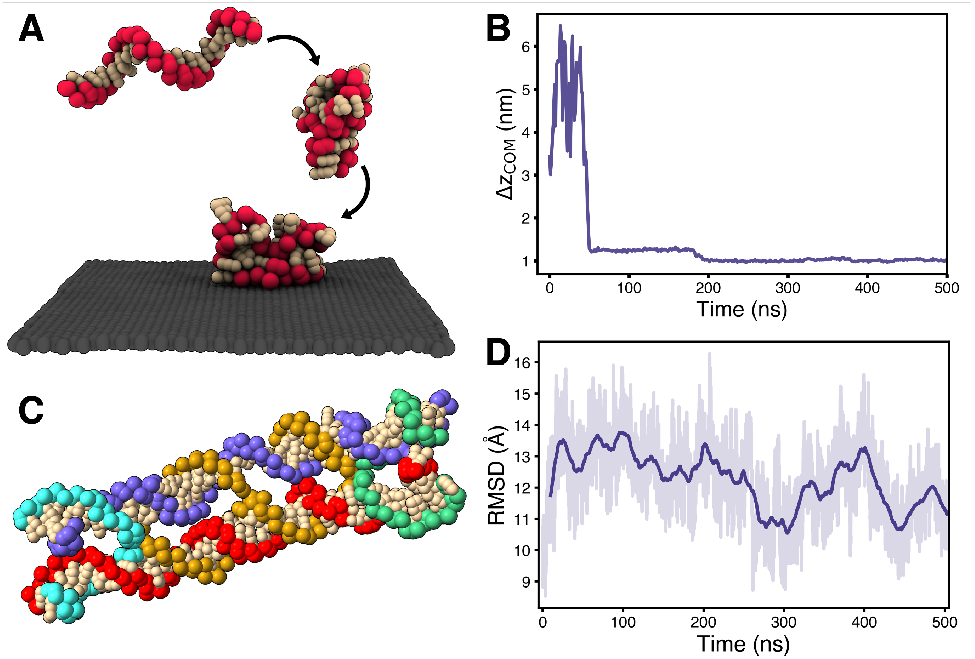
Application of the Martini 3 DNA model to graphene adsorption and DNA nanostructures. (A) Snapshots of ssDNA adsorbing onto a graphene sheet (dark gray). (B) Center-of-mass separation between the ssDNA and graphene surface along the sheet-normal, showing spontaneous adsorption and persistent surface association. (C) Snapshot of crossover-containing DNA nanostructure after approximately 500 ns of simulation. Each backbone strand is individually colored. (D) RMSD of the crossover-containing DNA nanostructure relative to the average structure. The light trace shows the instantaneous RMSD and the dark trace shows a 20 ns rolling average.

Finally, we tested the model in a crossover-containing DNA nanostructure consisting of two adjacent dsDNA duplexes connected by two Holliday junction crossovers [6, 7]. The structure remained intact over the 500 ns simulation, with no disruption of the crossover junctions or separation of the duplexes. The RMSD remained stable without a sustained increase over the trajectory (Fig. 7C,D).

Taken together, these applications show that the Martini 3 DNA model can be transferred beyond isolated DNA systems while retaining stable DNA structures and supporting the distinct interactions required by each environment. Stable protein binding, cholesterol-mediated membrane association, spontaneous ssDNA adsorption to graphene, and preservation of the crossover-containing nanostructure were all observed without reparameterizing the DNA model for each system. This ability to combine a single DNA representation with Martini 3 proteins, lipids, and non-biological materials supports its broader use in heterogeneous molecular simulations.

## Discussion

The optimized Martini 3 DNA model reproduced the local and global structural properties used during parameterization and showed improved agreement relative to the Martini 2 DNA model across these observables. Importantly, the model also reproduced structural and mechanical properties that were not explicitly included in the optimization objective, including B-DNA-like helicity and persistence lengths on the expected tens-of-nanometers scale. This suggests that the combination of local backbone and global duplex observables provided a sufficiently broad structural target for optimization. More generally, the results demonstrate the utility of the Bayesian optimization workflow as an effective strategy for coarse-grained force-field development, where multiple coupled structural properties must be balanced simultaneously rather than optimized independently.

A central motivation for extending DNA to Martini 3 was to enable simulations within the broader chemical space available in the Martini 3 framework. The application simulations demonstrate this across several distinct environments. The model maintained a stable *λ* repressor-DNA complex, preserved duplex integrity when cholesterol-tethered to a POPC membrane, spontaneously adsorbed to an independently parameterized graphene surface, and maintained a crossover-containing DNA nanostructure. These results show that the model can be combined with existing Martini representations of proteins, lipids, and non-biological materials in heterogeneous hybrid systems. This substantially broadens the range of DNA-containing systems accessible compared with the primarily biomolecular-focused scope of the Martini 2 force field [14, 15].

The current model nevertheless retains limitations that provide clear directions for further refinement. As in the Martini 2 DNA model, duplex stability relies on a fixed elastic network [14], which constrains the relative organization of the two strands and does not provide a mechanism for dynamic formation or disruption of base pairing. Consequently, the model cannot currently represent processes such as spontaneous duplex melting, hybridization, or strand exchange. Persistence length also showed little dependence on salt concentration, and the estimate at 500 mM remained above the commonly expected experimental range, although with substantial uncertainty. The overall magnitude and weak salt dependence were similar to those previously reported for Martini 2. The ssDNA simulations additionally produced structures that were more compact than experimental measurements, particularly for longer strands. Future refinement could therefore focus on enabling dynamic duplex formation and dissociation, improving the ion dependence of dsDNA mechanics, and incorporating ssDNA structural properties more directly into the parameterization. DNA nanotechnology represents a particularly promising application of the model. The stability of the crossover-containing structure provides an initial indication that Martini 3 DNA can support architectures beyond isolated duplexes. Larger DNA origami systems will provide more stringent tests of crossover mechanics, long-range structural stability, and accumulated strain. In addition, compatibility with the broader Martini 3 framework enables these nanostructures to be studied together with membranes, proteins, polymers, and other materials within a common coarse-grained representation.

## Acknowledgments

This work was supported by National Institutes of Health (Grant no. 2R01GM122979) and National Science Foundation (Grant no. CMMI-2323969). Computational resources were provided by the Duke Computing Cluster and the ACCESS program supported by the National Science Foundation (Grants nos. ACI-2138259, 2138286, 2138307, 2137603, and 2138296). The authors wish to acknowledge the contributions of Daniel Duke, for providing structural input on the DNA nanostructure, and Lisa Hall, for critical feedback on early versions of the model. AI tools were used to assist with manuscript refinement.

