## Supplemental Information for "Martini 3 Coarse-Grained Model of DNA for Heterogeneous Molecular Systems"

### Table of Contents

#### Supplemental Figures

- Figure 1: Distribution of observables sampled from optimized parameter set.
- Figure 2: $W_s$ and $D_{\mathrm{KL}}$ comparison across local observables.
- Figure 3: ssDNA simulation snapshots at various lengths and NaCl concentrations.
- Figure 4: Cholesterol-tagged DNA starting structure.
- Figure 5: Interactions between graphene and ssDNA.

#### Supplemental Tables

- Table 1: Bounds of optimization parameters.
- Table 2: Variations in CG simulation protocol across studied systems.
- Table 3: Simulated DNA sequences.

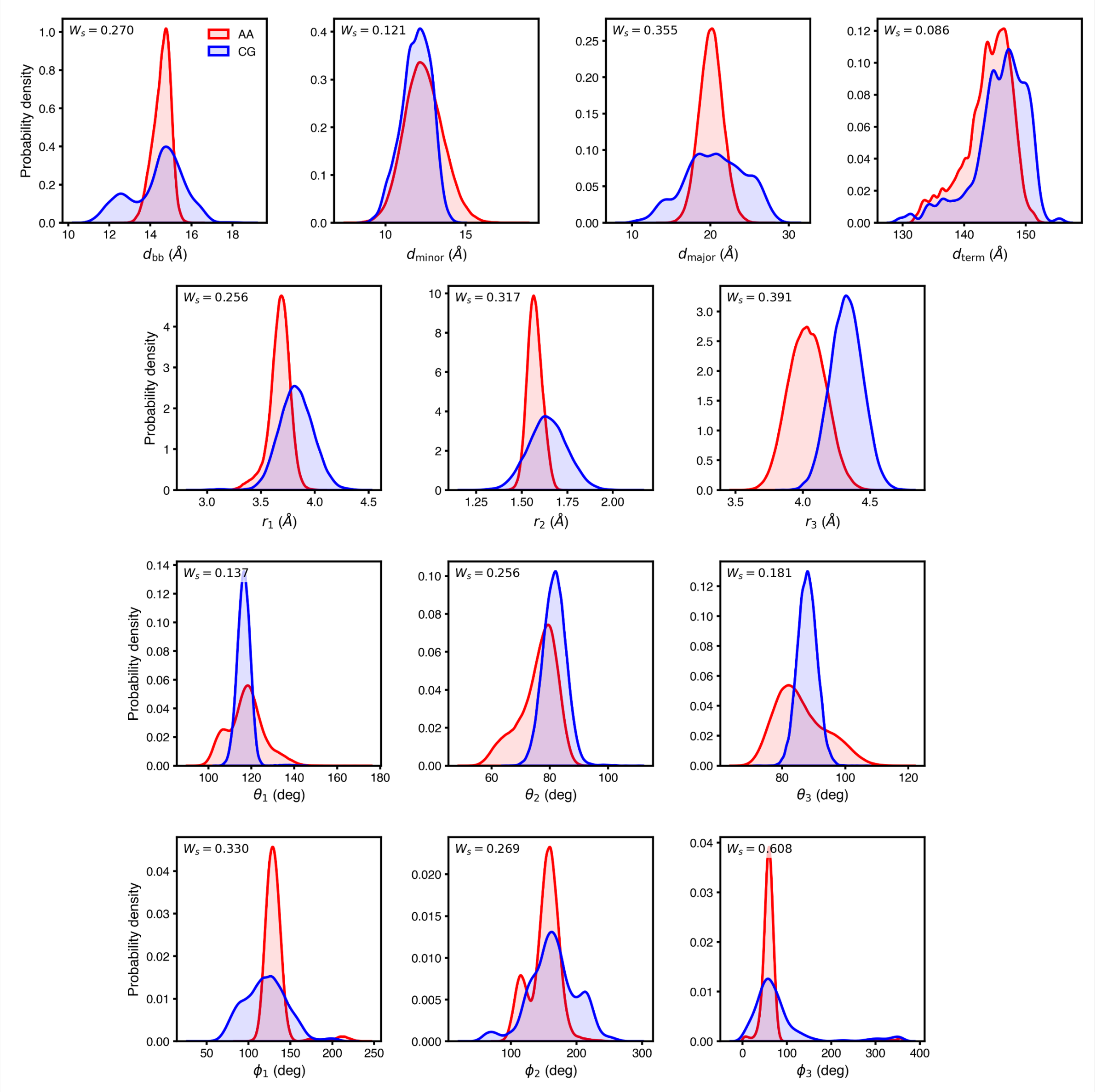


Figure 1: Distributions of observables for the optimized CG parameter set (blue) and AA reference simulations (red). The top row shows the four global observables used in the optimization, while the lower $3 \times 3$ grid shows the local backbone observables. The corresponding $W_s$ value is shown in the upper-left corner of each panel.

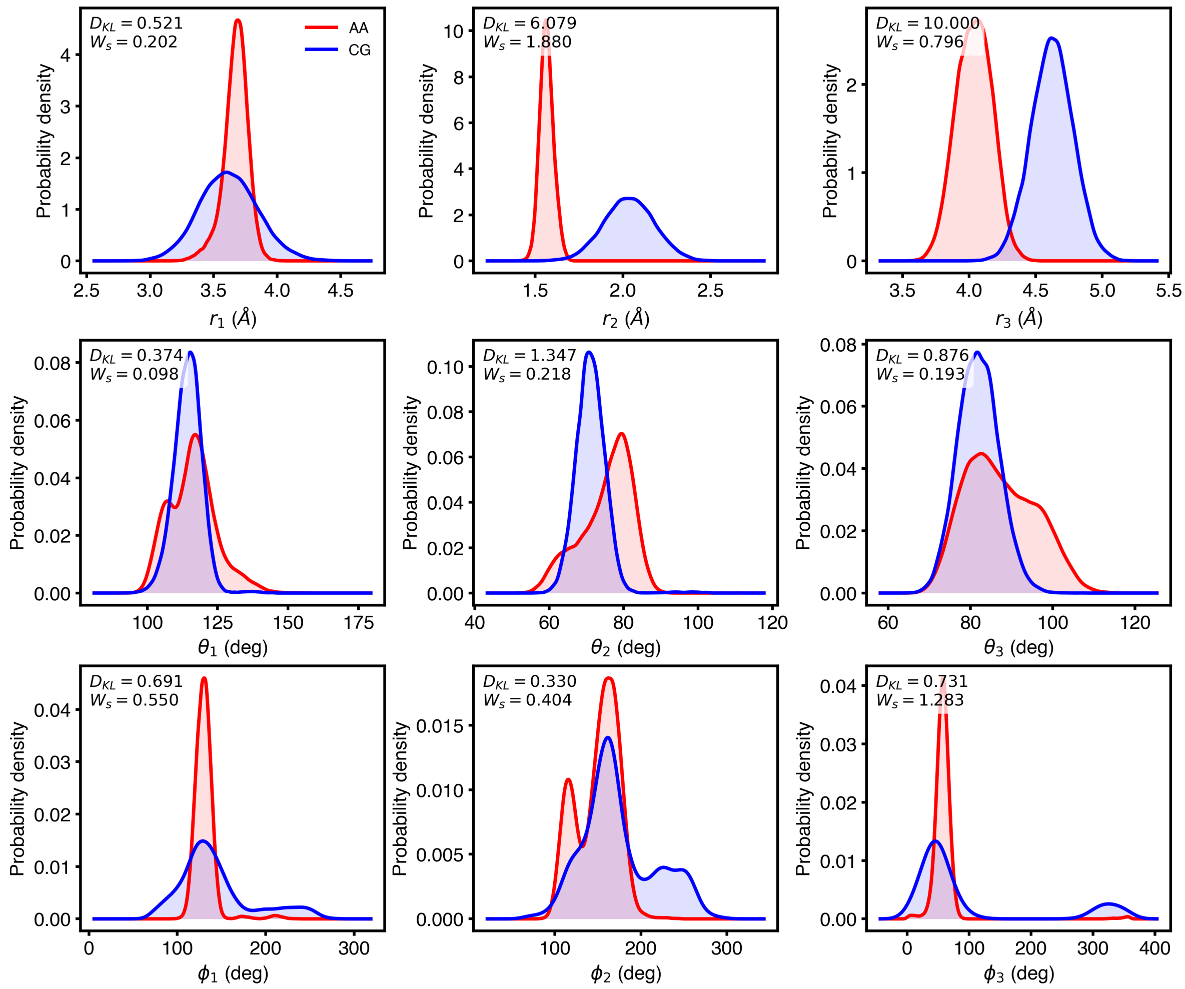


Figure 2: $W_s$ and $D_{\mathrm{KL}}$ for local observables from a randomly selected parameter set. Calculated $W_s$ and $D_{\mathrm{KL}}$ values are shown in the upper-left corner of each panel. The $r_3$ distributions show greater visual agreement between AA and CG than $r_2$, consistent with the lower $W_s$ for $r_3$, whereas $D_{\mathrm{KL}}$ assigns a larger value to $r_3$. This is one of several cases in which $W_s$ was more consistent with visual agreement between the AA and CG distributions than $D_{\mathrm{KL}}$.

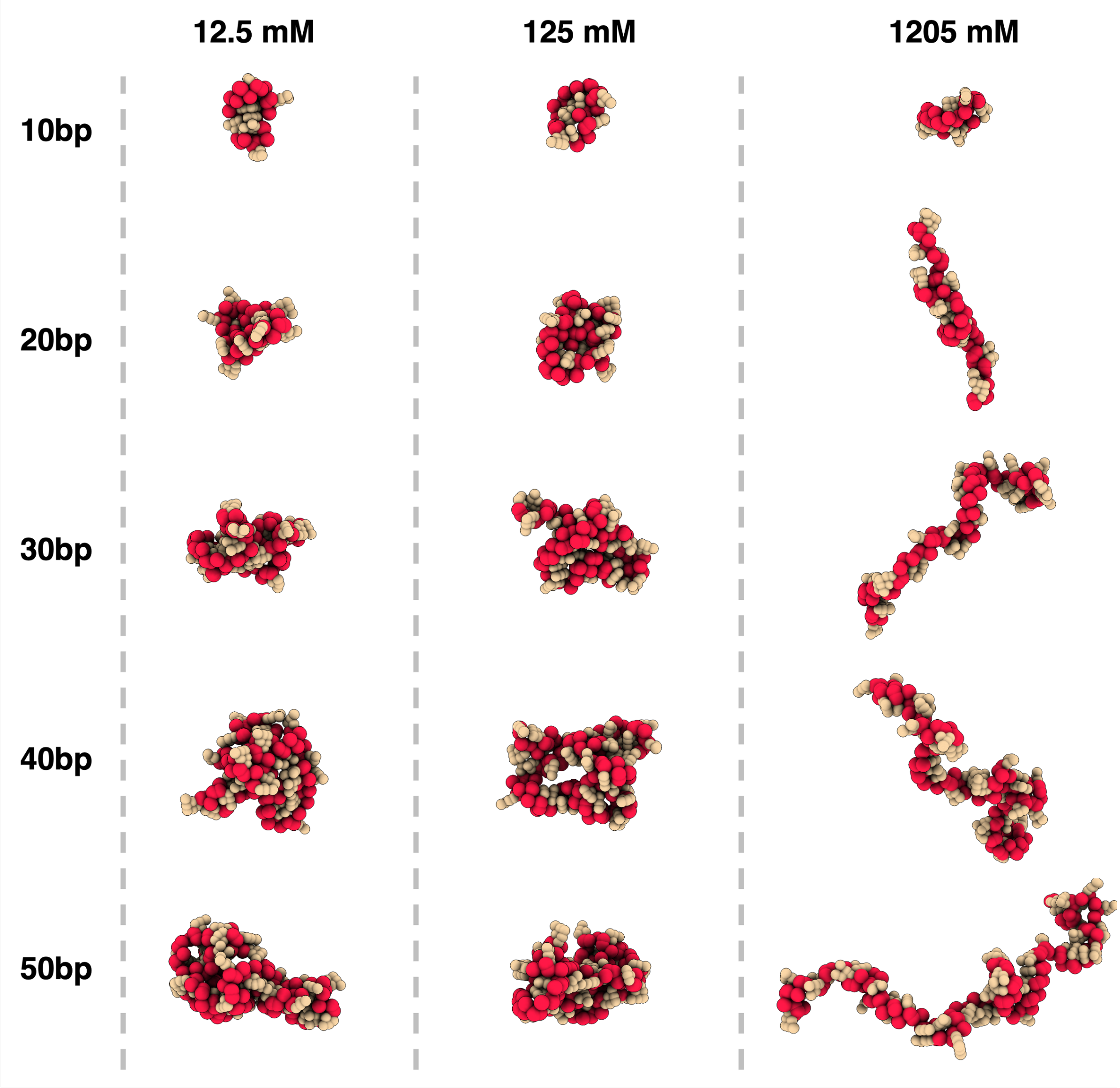


Figure 3: Representative snapshots of ssDNA used for radius-of-gyration calculations after 80 ns of production simulation. Rows correspond to ssDNA lengths of 10, 20, 30, 40, and 50 nt, and columns correspond to NaCl concentrations of 12.5, 125, and 1205 mM. At 1205 mM NaCl, the ssDNA remained comparatively extended rather than exhibiting the compaction observed at the lower salt concentrations. This non-monotonic behavior may reflect altered ion-DNA and ion-solvent interactions at very high ionic strength, substantially above the physiological salt concentrations commonly used in Martini 3 simulations.

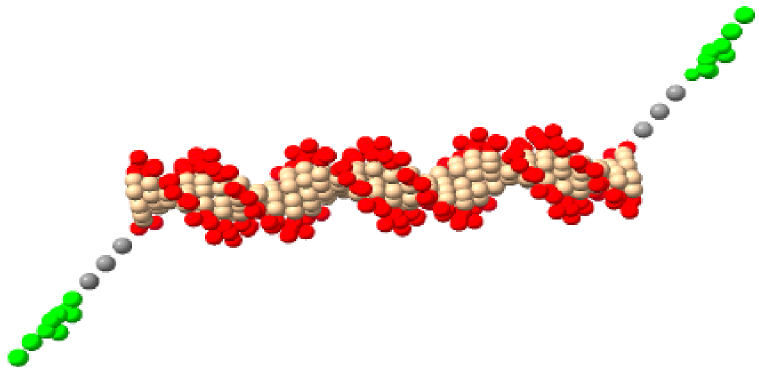

Figure 4: Starting CG structure of cholesterol-tagged dsDNA. Cholesterol (green beads) was attached to the DNA (red and tan beads) through a six-carbon aliphatic linker (gray beads), coarse-grained into three beads with types SC3-SC3-SP4. The terminal SC3 bead was attached to the 5′ end of the DNA at the BB2 bead, while the SP4 bead was attached to the cholesterol ROH bead.

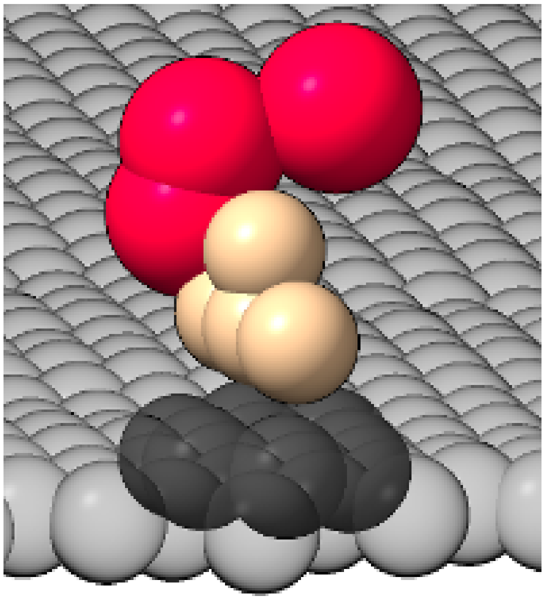

Figure 5: Zoomed-in snapshot of a T-shaped $\pi$-$\pi$ stacking interaction between ssDNA and graphene during the CG simulation. A single interacting nucleotide from the longer ssDNA strand is shown for clarity, while the remainder of the strand is hidden. The ssDNA backbone is shown in red and the nucleobase in tan. The graphene sheet is shown in dark gray, with the interacting aromatic ring rendered opaque and the surrounding sheet transparent.

Table 1: Lower and upper bounds used for optimization of the Martini 3 DNA backbone bonded parameters.

| **Parameter** | **Lower bound** | **Upper bound** |
|---|---|---|
| $r_1^0$ | 0.25 nm | 0.45 nm |
| $k_{r_1}$ | 800 kJ/(mol nm$^2$) | 24,000 kJ/(mol nm$^2$) |
| $r_2^0$ | 0.10 nm | 0.20 nm |
| $k_{r_2}$ | 800 kJ/(mol nm$^2$) | 24,000 kJ/(mol nm$^2$) |
| $r_3^0$ | 0.30 nm | 0.50 nm |
| $k_{r_3}$ | 800 kJ/(mol nm$^2$) | 20,000 kJ/(mol nm$^2$) |
| $\theta_1^0$ | 90° | 150° |
| $k_{\theta_1}$ | 100 kJ/mol | 1,000 kJ/mol |
| $\theta_2^0$ | 50° | 110° |
| $k_{\theta_2}$ | 100 kJ/mol | 1,000 kJ/mol |
| $\theta_3^0$ | 60° | 120° |
| $k_{\theta_3}$ | 50 kJ/mol | 1,000 kJ/mol |
| $\phi_1^0$ | 90° | 180° |
| $k_{\phi_1}$ | 0 kJ/mol | 50 kJ/mol |
| $\phi_2^0$ | 100° | 200° |
| $k_{\phi_2}$ | 0 kJ/mol | 50 kJ/mol |
| $\phi_3^0$ | 0° | 100° |
| $k_{\phi_3}$ | 0 kJ/mol | 50 kJ/mol |

Table 2: Simulation conditions and protocol variations for the CG systems.

| Sim. Name | NaCl* | Length | Protocol Variation** |
|---|---|---|---|
| Parameter optimization (dsDNA) | 150 mM | 100 ns | None |
| Helicity and persistence length (dsDNA) | 0, 150, 500 mM | 500 ns | 300 K |
| Radius of gyration (ssDNA) | 12.5, 125, 1205 mM | 100 ns | None |
| dsDNA-CHOL on POPC Bilayer | 150 mM | 500 ns | 2 fs timestep; semi-isotropic pressure coupling |
| ssDNA on graphene sheet | 150 mM | 1000 ns | Tiny water beads; graphene position restrained |
| $\lambda$ repressor-operator complex (dsDNA) | 150 mM | 625 ns | 2 fs timestep |
| DNA nanostructure | 150 mM | 500 ns | 2 fs timestep |

*NaCl concentration after neutralization with $Na^+$ counterions.

**Protocol variations relative to the default CG simulation protocol described in the Methods.

Table 3: DNA sequences used in the simulations.

| Sim. Name | Length | Sequence (5′-3′) |
|---|---|---|
| Parameter optimization (dsDNA) | 50 bp | AAGTTTAACA AATTATCTCG ATCGGTTGAA CAGGGACACA GGTAATGAGG |
| Helicity and persistence length (dsDNA) | 100 bp | GTTAACGGCA GGTGCAACCC ATTGTTGCAG CGTAGGCACC GTCGCTTGCC CTCGTGGCAC GGGCGTCGAT GCAGGATTCA TCGGCTTCGC TGTTCGCTGT |
| Radius of gyration (ssDNA) | | |
| | 10 nt | TATCGGTATG |
| | 20 nt | GCTTACTAGC TCGTGGTTGT |
| | 30 nt | ATGTGGCATA AGATTTATTG GCGCGACGGC |
| | 40 nt | CCGCATACGT TTGCTACGAG TAACAGACCT AATTCCGGGC |
| | 50 nt | ATGGTACAGA TCCTATCCTA CCCTATGCGC TGGACCTGAC CAATGAGGGT |
| dsDNA-CHOL on POPC Bilayer | 30 bp | GAACTCGTGG CTTGGACATC AGAAATACCC |
| ssDNA on graphene sheet | 20 nt | GCTTACTAGC TCGTGGTTGT |
| $\lambda$ repressor-operator complex (dsDNA) | 20 bp | AATACCACTG GCGGTGATAT |
| DNA nanostructure* | | |
| Strand A | 41 nt | CTCTTTGTCC TGTCAGGGAC GTGTATAAGA AGAGGCGTCC C |
| Strand B | 41 nt | AATTCTGTCT ATCATATGGC TAAGGCCGAC AATATGGTCT A |
| Crossover strand 1 | 20 nt | GGGACGCCTC AGACAGAATT |
| Crossover strand 2 | 42 nt | CGTCCCTGAC ATGTCGGCCT TAGCCATATG ATTTCTTATA CA |
| Crossover strand 3 | 20 nt | TAGACCATAT GGACAAAGAG |

*The DNA nanostructure consists of two 41 bp duplexes connected by two crossover regions. Crossovers occur between residues 10 and 11 of crossover strands 1 and 3, and between residues 11 and 12 and residues 32 and 33 of crossover strand 2.
